# A 3.4-Å cryo-EM structure of the human coronavirus spike trimer computationally derived from vitrified NL63 virus particles

**DOI:** 10.1101/2020.08.11.245696

**Authors:** Kaiming Zhang, Shanshan Li, Grigore Pintilie, David Chmielewski, Michael F. Schmid, Graham Simmons, Jing Jin, Wah Chiu

**Affiliations:** Department of Bioengineering and James H. Clark Center, Stanford University, Stanford, CA 94305, USA; Graduate Program in Biophysics, Stanford University, Stanford, CA 94305, USA; Vitalant Research Institute, San Francisco, CA 94030, USA; Division of CryoEM and Bioimaging, SSRL, SLAC National Accelerator Laboratory, Menlo Park, CA 94025, USA

**Keywords:** Human coronavirus NL63, spike trimer, cryo-EM

## Abstract

Human coronavirus NL63 (HCoV-NL63) is an enveloped pathogen of the family *Coronaviridae* that spreads worldwide and causes up to 10% of all annual respiratory diseases. HCoV-NL63 is typically associated with mild upper respiratory symptoms in children, elderly and immunocompromised individuals. It has also been shown to cause severe lower respiratory illness. NL63 shares ACE2 as a receptor for viral entry with SARS-CoV and SARS-CoV-2. Here we present the *in situ* structure of HCoV-NL63 spike (S) trimer at 3.4-Å resolution by single-particle cryo-EM imaging of vitrified virions without chemical fixative. It is structurally homologous to that obtained previously from the biochemically purified ectodomain of HCoV-NL63 S trimer, which displays a 3-fold symmetric trimer in a single conformation. In addition to previously proposed and observed glycosylation sites, our map shows density at other amino acid positions as well as differences in glycan structures. The domain arrangement within a protomer is strikingly different from that of the SARS-CoV-2 S and may explain their different requirements for activating binding to the receptor. This structure provides the basis for future studies of spike proteins with receptors, antibodies, or drugs, in the native state of the coronavirus particles.

## Introduction

*Coronaviridae* constitute a large family of enveloped, positive-sense single-stranded RNA (+ssRNA) viruses capable of causing severe and widespread human respiratory disease. The coronaviruses are zoonotic pathogens, often circulating among natural reservoirs such as bats or camels prior to crossing the species barrier into humans (Sabir *et al.* 2016; Wang *et al.* 2006). Coronaviruses are classified into four genera (alpha-CoV, beta-CoV, gamma-CoV and delta-CoV), with human coronaviruses found in two: alpha-CoVs (HCoV-229E and HCoV-NL63) and beta-CoVs (middle-east respiratory syndrome (MERS) and severe acute respiratory syndrome coronavirus 1 and 2 (SARS-CoV-1, SARS-CoV-2)) (Cui *et al.* 2019). SARS-CoV-2, responsible for the COVID-19 pandemic, has infected over 19 million people and claimed over 720,000 lives worldwide as of early August, 2020, underscoring the urgency of studying all circulating human coronaviruses (https://covid19.who.int/) (Cui *et al.* 2019; Zhou *et al.* 2020). While beta-CoVs are more commonly associated with high pathogenesis and severe respiratory disease, alpha-CoVs are widely circulating viruses associated with cold-like symptoms and in rare cases more serious respiratory failure (Wang *et al.* 2020). HCoV-NL63 is estimated to be the causative agent of up to 10% of annual respiratory disease and is a major cause of bronchiolitis and pneumonia in newborns (Chiu *et al.* 2005; van der Hoek *et al.* 2004).

Coronaviruses utilize large (~500-600 kDa) spike (S) homotrimers protruding from the viral membrane to engage cellular receptors and mediate fusion with host membranes (Pallesen *et al.* 2017; Thorp *et al.* 2006). These spikes, numerous on the virus surface, constitute the primary target of neutralizing antibodies (NAbs) and are central to vaccine design and structural studies for drug optimization. Each S molecule contains two large regions: N-terminal S1 responsible for receptor-binding and C-terminal S2 responsible for type-I fusion, as well as an additional single-pass transmembrane helix that anchors the spikes to the viral envelope (Bosch *et al.* 2003; Zheng *et al.* 2006). Spikes are activated by protease cleavage at the S2’ site, near the putative fusion peptide, allowing the transition from pre-to post-fusion conformation during virus entry (Belouzard *et al.* 2009).

Several single-particle cryo-electron microscopy (cryo-EM) structures of purified, ectodomains of pre-fusion S trimers from alpha-CoV and beta-CoV genera have been determined, illustrating conformational heterogeneity of S1, multiple glycosylation sites, and the interactions with soluble receptors and NAbs (Song *et al.* 2018; Walls *et al.* 2016, 2020; Wrapp *et al.* 2020). These S complexes lack the helical C-terminal stem that connects S2 to the viral envelope, and require residues mutated to proline at the loop between the first heptad repeat (HR) and the central helix to stabilize the construct (Kirchdoerfer *et al.* 2018; Pallesen *et al.* 2017; Park *et al.* 2019). High resolution structure determinations of molecular components in pleomorphic virions are typically obtained by single particle cryo-EM or X-ray crystallography of exogenously expressed and purified components. Conversely, cryo-electron tomography (cryo-ET) has been used to reconstruct tomograms of the whole virus particles, from which subvolume density, corresponding to the feature of interest, would be further processed by classification and averaging (Schmid *et al.* 2012; Wan & Briggs 2016). Here, we take the approach of utilizing single-particle cryo-EM imaging of purified and vitrified HCoV-NL63 virus particles, and then using computational methods to extract the S trimer density for its high resolution structure determination. Our approach is similar to a recent preprint report for chemically fixed SARS-CoV-2 particles (Ke *et al.* 2020). We report the 3.4-Å structure of the S trimer in its native, pre-fusion state, and expect our data will be of value in future drug and vaccine design for various strains of coronaviruses without the need to genetically construct and biochemically purify spike proteins.

## Results and Discussion

Since the S timers protrude from the membrane surface of the virion, many of the crown-shaped S trimers can be easily identified, computationally extracted from the raw images, and treated as single particle images with a distinct particle orientation without the need of tilting the specimens (**Fig. 1a**). Based on the fact that single-particle cryo-EM is a more routine procedure for computing high-resolution structures compared to cryo-ET, we decided not to collect tilt series, and used single-particle cryo-EM analysis of the native S trimer on purified and vitrified virions. The 2D class averages showed a clear triangular shape for the S trimer and other distinct views (**Fig. 1b**). D*e novo* building of the initial map, using the “Ab initial reconstruction” option in cryoSPARC without any symmetry applied, resulted in a three-dimensional (3D) structure with well-defined features. Further 3D refinement was performed with and without C3 symmetry to obtain 3.4 Å and 3.7 Å maps, respectively (**Fig. 1c-e and Fig. S1**). The reconstructed map without imposed symmetry also displayed 3-fold symmetry, showing a high cross-correlation coefficient (CC = 0.9802) with the symmetric map (**Fig. S1d**). Hereafter, we will present our structural analysis of the 3.4-Å map with C3 imposed symmetry after refinement from a subset of ~82,000 S trimer particles showing adequate orientation sampling (**Fig. S1**). The local resolution varies in our map (**Fig. 1e**); the densities at the utmost surface of the protein have a much lower resolution than the central region (**Fig. 1e**), which could be attributed to the inherent flexibility of the distal ends of glycans. Compared to the map of the purified S ectodomain, our map has more extended density without recognizable secondary structure element features pointing toward the viral membrane (**Fig. 1e**). The detection of this feature likely benefits from the direct picking of S trimer anchored in the lipid envelope of native virions, stabilizing the stem region that otherwise disassembles in the purified S ectodomains.

**Figure 1.**
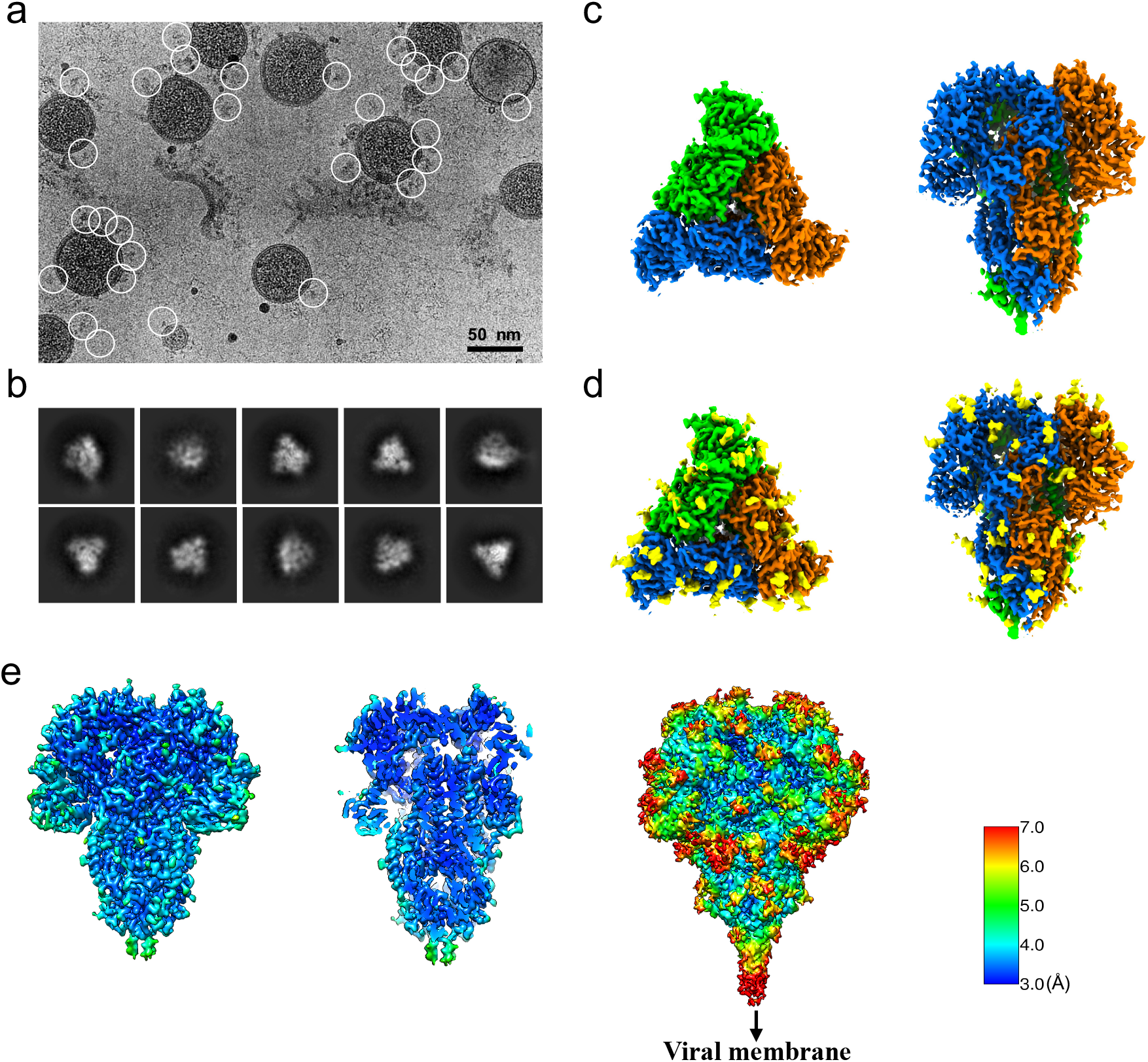
Single particle cryo-EM analysis of *in-situ* structure of the HCoV-NL63 coronavirus spike glycoprotein. (a) Representative motion-corrected cryo-EM micrograph. (b) Reference-free 2D class averages of computationally extracted spikes. (c-d) Reconstructed cryo-EM map of the spike in top and side views without (c) and with (d) glycans shown. (e) Resolution variation maps for the 3D reconstruction. Left, whole map view; middle, slice view; right, whole map view at a lower threshold.

Because of the high feature similarity of our map and that determined from the biochemically purified S ectodomain (Walls *et al.* 2016), we fit the published model (PDB ID: 5SZS) to our 3.4-Å map and refined it including the glycans (see Methods). The quality of the refined model was validated by MolProbity (Chen *et al.* 2010) (**Table S1**), correlation between the map and model, and Q-score analysis per residue (Pintilie *et al.* 2020) (**Fig. 2**). The density map of our structure is better resolved at many regions relative to the previous structure (PDB ID: 5SZS), including glycans (**Fig. 3**), which were previously interpreted with accompanied mass spectroscopy measurements (Walls *et al.* 2016). Superimposition of our model and 5SZS yields 804 pruned atom pairs matched with 1.28-Å RMSD, indicating their high structural similarity. Structural comparisons based on individual domains also show similar results. These domains include domain 0, domain A, domain B (also known as RBD), domain C, and domain D, with the RMSD ranging from 0.29 Å to 0.90 Å. As shown in **Fig. 1 and 2**, each of the three receptor-binding domains (RBDs) within the S trimer is pointed downwards. Notably, heterogeneous refinement and 3D variability did not reveal alternative RBD conformations, implying that the native S trimer proteins on the virion are predominantly in a fully closed state, consistent with the previous report of purified HCoV-NL63 S ectodomain (Walls *et al.* 2016).

**Figure 2.**
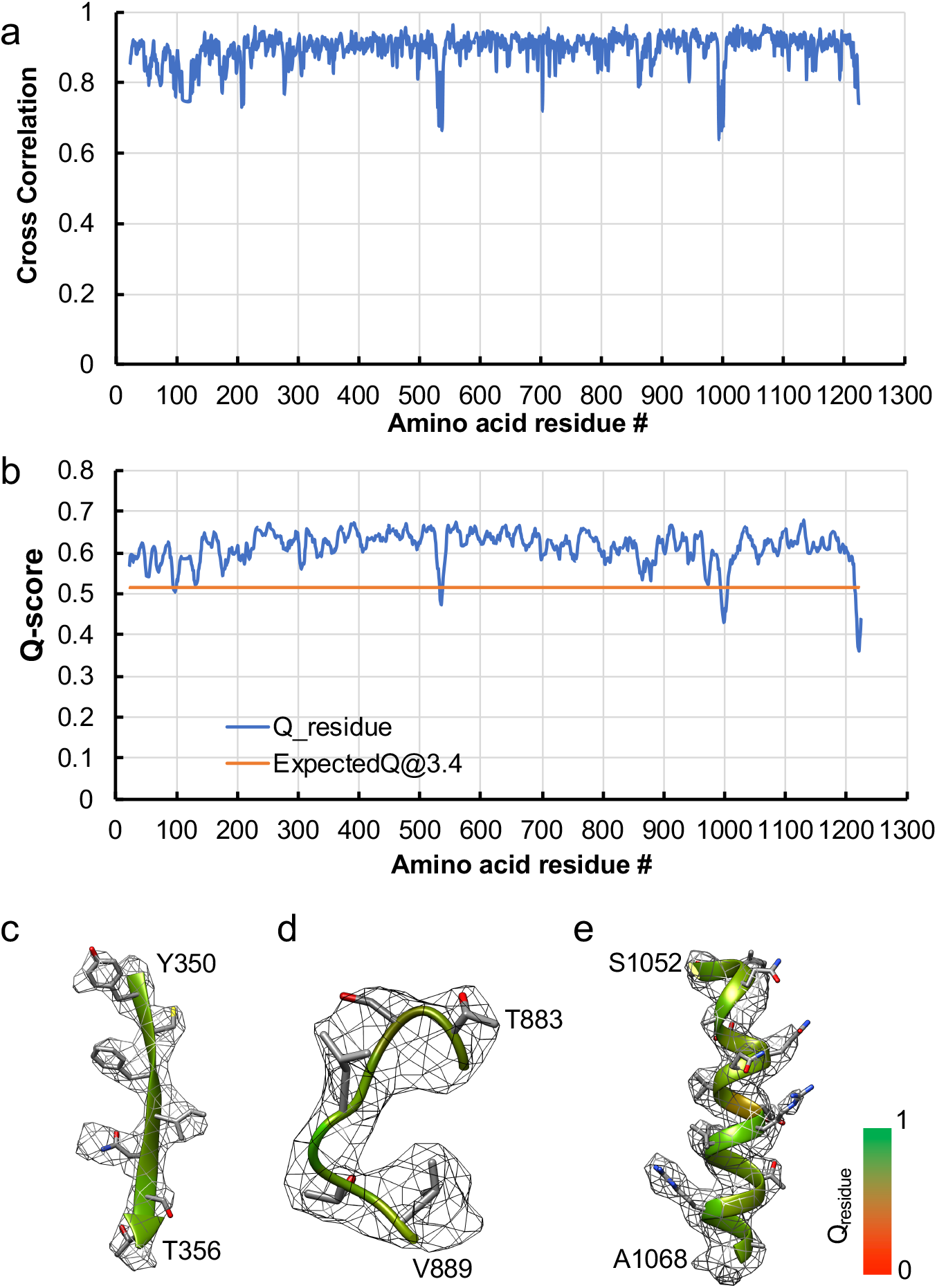
Model validation of the HCoV-NL63 coronavirus spike glycoprotein protomer. (a) Per-residue cross-correlation coefficient between the model and 3.4-Å map. (b) Q-score for each amino acid residue in model and 3.4-Å map; the orange line represents the expected Q-score at 3.4-Å resolution (0.5157) based on the correlation between Q-scores and map resolution (Pintilie et al. 2020). (c-e) Sample density maps of various regions. The model is shown as ribbon, with residue Q-scores indicated. The higher Q-score indicates better resolvability. (c) β-sheet. (d) Residues were not built in the biochemically purified HCoV-NL63 spike protein structure (PDB ID: 5SZS). (e) α-helix.

**Figure 3.**
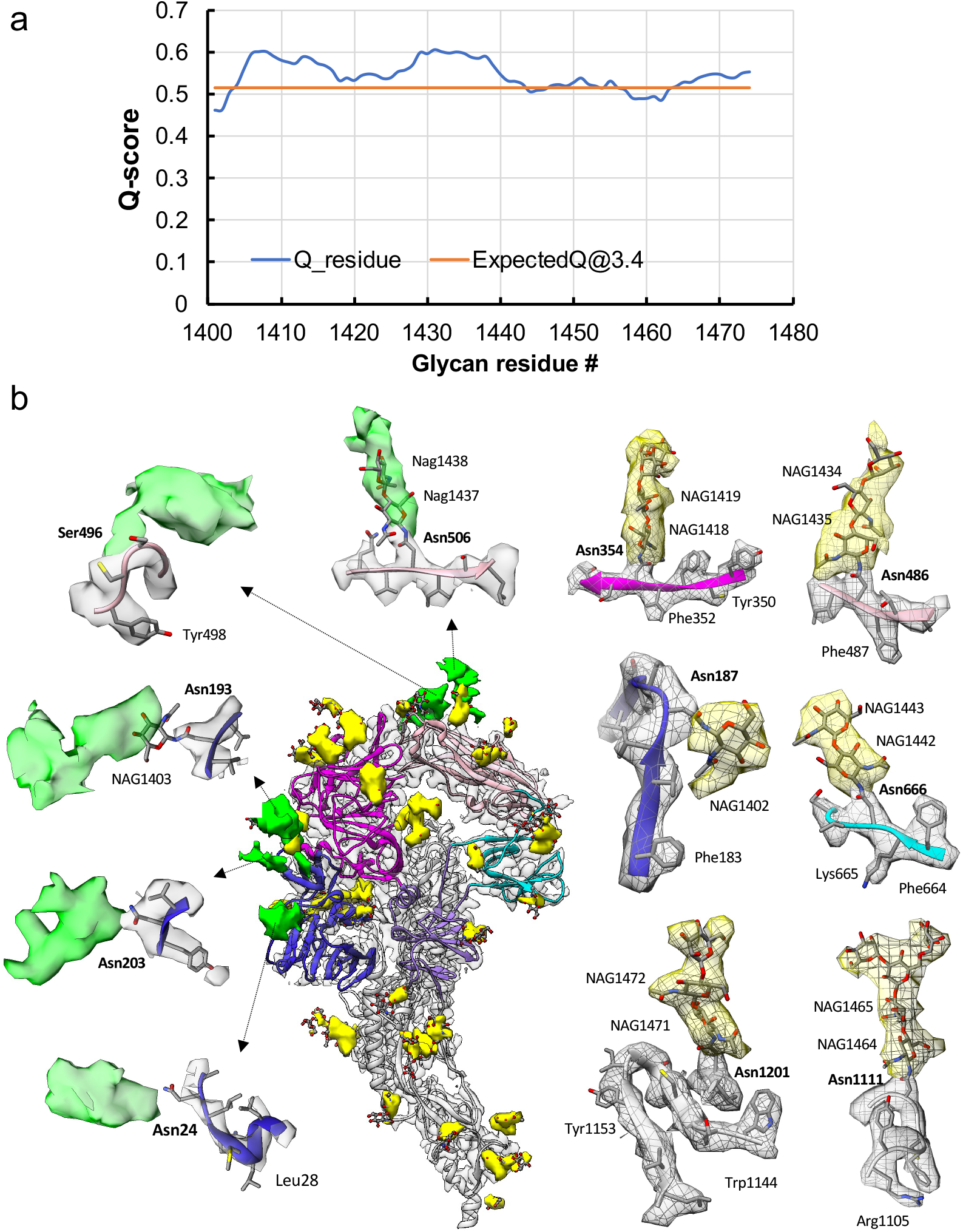
Resolvability of glycans. (a) Q-scores analysis for each glycan residue (# starting from 1401) in model and map; the orange line represents the expected Q-score at 3.4-Å resolution (0.5157) based on the correlation between Q-scores and map resolution of amino acid residues (Pintilie et al., 2020). (b) Highlights on several glycans. Green: possible glycan densities in our map compared to previous study (EMD-8331); Yellow: glycan densities derived from the model in previous study (PDB ID: 5SZS).

Since glycosylation of S plays an important role in the viral life cycle and immune-evasion (Casalino *et al.* 2020; Vigerust & Shepherd 2007; Watanabe *et al.* 2019, 2020), we further investigated the glycosylation sites in our map by computing a difference map between our map and the map of biochemically purified S (EMD-8331). All glycosylation sites previously identified by a combination of cryo-EM densities protruding from the expected amino acid side chains and mass spectroscopy (**Table 1 and Table S2**) (Walls *et al.* 2016) were found in our map and annotated in yellow (**Fig. 3**) as in (EMD-8331). Two other N-linked glycosylation sites (Asn24 and Asn203 in domain 0), predicted to exist from their sequence but not found by Mass Spectroscopy or in the map of (Walls *et al.* 2016), as well as one O-linked glycosylation site (Ser496 in domain B), were newly identified in our density map (**Fig. 3, Table 1**). In addition, we noted extended densities at several glycosylation sites shown in the difference map (annotated in green, **Fig. 3**) between our structure and the previously published one, possibly due to the differences in glycosylation patterns across species (Walls *et al.* 2016). In our study the virus was grown in mammalian cells, while the previous study used S ectodomain expressed in insect cells. N-linked glycosylation in mammalian cells typically results in complex-type glycans with two to four branches extended from the tri-mannosyl core. In contrast, glycosylation in insect cells typically yields truncated or paucimannosidic or oligomannosidic glycans, with few if any complex-type glycans (Tomiya *et al.* 2004). Overall there is significant agreement between *in-situ* and biochemically purified S protein structure, with some notable differences in glycan densities. Our structure cross-validates the conformation of the spike protein of HCoV-NL63 determined from either the biochemically purified specimen or the intact virions by cryo-EM.

**Table 1.**

| Site | Sequon | MS identified<br>(Walls et al) | Cryo-EM<br>observed density<br>(Walls et al) | Cryo-EM observed<br>density (this study) |
| --- | --- | --- | --- | --- |
| 24 | <u>N</u>LSM | ND | ND | 24 |
| 203 | <u>N</u>YTV | ND | ND | 203 |
| 496 | GG<u>S</u>C | ND | ND | 496 |

Compared to the S1 protein of betacoronavirus (i.e. SARS-CoV-2), HCoV-NL63 S1 has an additional N-terminal domain (domain 0) that is assumed to be a gene duplication of domain A and is a canonical feature of alphacoronaviruses. Domain B (RBD) of SARS-CoV-2 transitions between upward (open) and downward (closed) conformations due to its inherent flexibility, the viral receptor ACE2 is able to sample the RBD’s up conformation to initiate virus-receptor binding. In contrast, domain B of HCoV-NL63 interacts with domain A, therefore being stabilized in a closed “circle” of HCoV-NL63 S1 (**Fig. 4a**). The interface area between domain A and B is ~500 Å^2^ and involves 22 hydrophobic interactions (**Fig. 4b**) according to PDBsum structure bioinformatics analysis (Laskowski *et al.* 2018). The three receptor binding motifs (RBMs) on HCoV-NL63 RBD are consequently buried in the interface between domain A and domain B, and prevented from binding to the viral receptor ACE2 (Wu *et al.* 2009). There must exist a mechanism to induce some conformational change to release the RBMs from domain A to allow receptor binding. It was reported that HCoV-NL63 utilizes heparan sulfate proteoglycans as the attachment factor before binding to ACE2 for virus entry (Milewska *et al.* 2014, 2018). Whether heparan sulfate binding to HCoV-NL63 S1 or other factors induce conformational changes to release RBMs for ACE2 binding warrants further studies.

**Figure 4.**
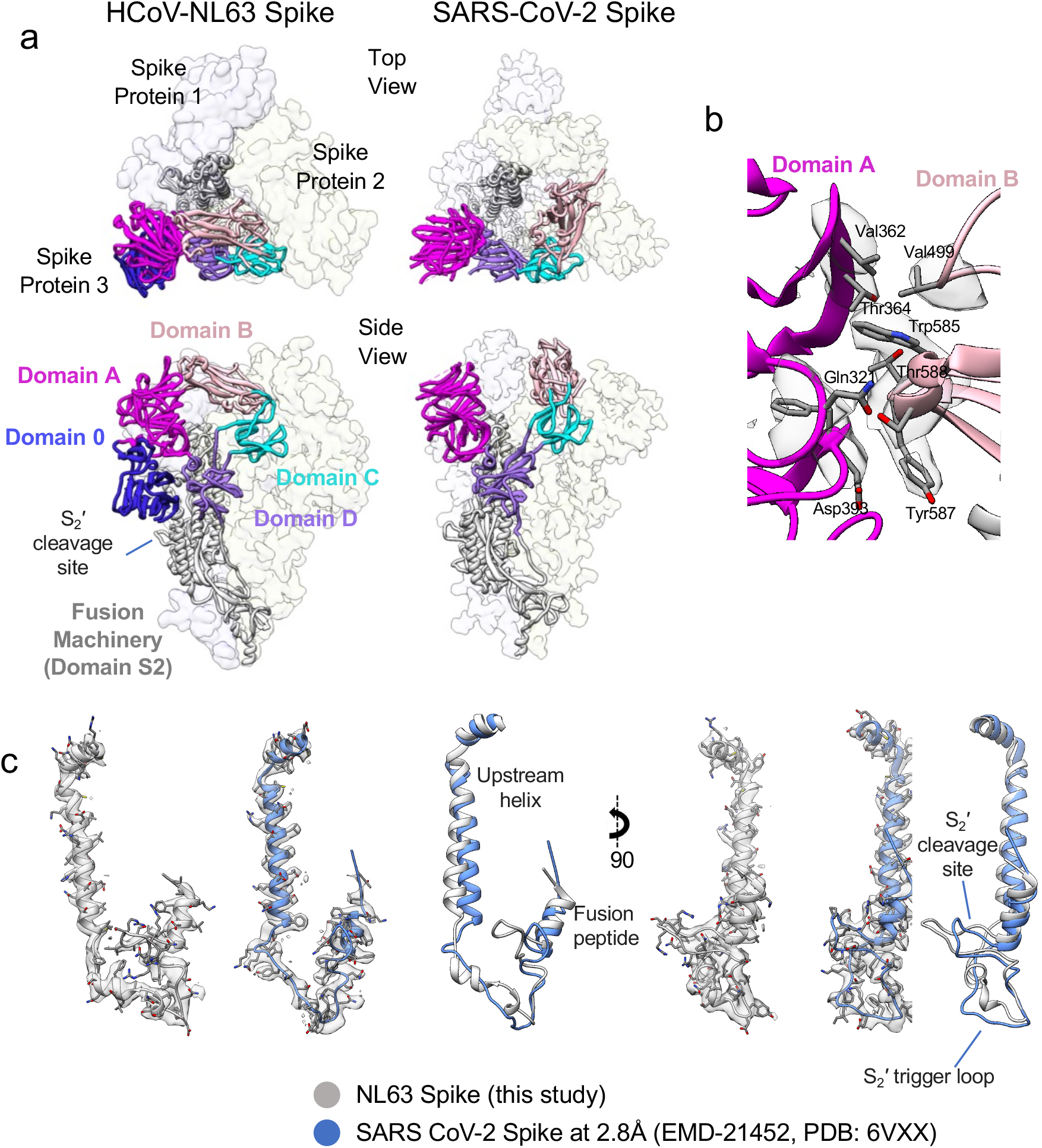
Map comparison between our structure and SARS-CoV-2 spike. (a) Comparison of our structure with SARS-CoV-2 spike in closed state (PDB ID: 6VXX) in the top view and side view. Each domain in the S1 subunit is shown in different colors. (b) Zoom-in view to show the interactions between domain A and domain B in the HCoV-NL63 spike protein. (c) Extracted densities of the S2’ trigger-loop region with models fitted in two views.

S proteins of coronaviruses require protease cleavage to release the fusion peptide that inserts into the target membrane to initiate membrane fusion for virus entry. We compared HCoV-NL63 and SARS-CoV-2 S2 fusion machineries by superimposing their structures together and excellent agreement was observed (**Fig. 4c**). In both structures, the S2’ loop where the host protease cleavage site is located is almost perpendicular to the central helix, protruding at the peripheral of the S trimer and readily accessible to host proteases. The fusion machineries of HCoV-NL63 and SARS-CoV-2 are almost identical except that the S2’ trigger loop of HCoV-NL63 forms a short alpha helix before looping back to connect to the fusion peptide while the corresponding loop of SARS-CoV-2 is long and flexible. The fusion machinery of coronaviruses provides a potential target for broad anti-coronavirus therapeutics development.

The feasibility of solving the near-atomic resolution structure of membrane-anchored S trimer on the purified viral particle opens a straightforward path for follow-up structural studies of coronaviruses with external reagents like receptors, antibodies or drugs. In addition, the single particle cryo-EM derived structures can be used together with subtomogram averages of the S trimer in the virion to understand complex and heterogeneous modes of interactions between S and various external reagents in different biochemical conditions, and the orientation in 3D of S with respect to the viral membrane, and, ultimately, the cell surface. Such a hybrid approach across different imaging protocols will be very useful for the discovery of authentic structural states of molecular components in a pleomorphic virus particle in the context of pathogenesis relevant to human health.

## Methods

### Cell culture and Virus

Vero E6 (ATCC CRL-1586) and MA104 (ATCC CRL-2378.1) cells were maintained at 37°C in a fully humidified atmosphere with 5% CO_2_ in DMEM (Invitrogen) and M199 (Gibco) medium respectively. All culture media were supplemented with penicillin and streptomycin and 10% FBS (Hyclone). HCoV-NL63 was obtained from BEI Resources, NIAID, NIH: Human Coronavirus, NL63, NR-470.

### Virus production and purification

Monolayers of Vero E6 or MA-104 cells were infected with HCoV-NL63 at MOI 0.5. Culture supernatants were harvested when clear cytopathic effect (CPE) developed. After clarification of the supernatants through a 0.45 micron filter, the virus was pelleted down through a 20% sucrose cushion. Virus was then resuspended in NTE buffer (20 mM Tris, pH 8.0, 120 mM NaCl, 1 mM EDTA) before layered onto 2 ml of 60% OptiPrep (Sigma-Aldrich) and spun at 50,000 x g for 1.5 hour using a SW28 rotor. After ultracentrifugation, the supernatants were removed to leave 4 ml above the virus band. The remaining 4 ml supernatant, the virus band and the underlay of 2 ml of 60% OptiPrep were mixed to reach a final concentration of 20% OptiPrep. The mixture was spun at 360,000 x g for 3.5 hours with a NVT65.2 rotor. The virus band was extracted and buffer exchanged to the NTE buffer using an Amicon Ultra-2 Centrifugal Filter Unit with Ultracel-100 membrane (Millipore).

### Cryo-EM vitrification, data acquisition, image processing, and 3D reconstruction

Three microliters of purified HCoV-NL63 virions (4.5 mg/mL) were first applied to 300-mesh R2/1 + 2 nm C-film grids (Quantifoil) that had been glow-discharged for 15s PELCO easiGlow (TED PELLA, INC.). Grids were then front-blotted for 2 seconds in a 90% humidified chamber and vitrified using the LEICA EM GP automated plunge freezing device. Frozen grids were then stored in liquid nitrogen until imaging. The samples were imaged in a Titan Krios cryo-electron microscope (Thermo Fisher Scientific) operated at 300 kV with GIF energy filter (BioQuantum, Gatan) at a magnification of 64,000× (corresponding to a calibrated sampling of 1.4 Å per pixel). Micrographs were recorded by EPU software (Thermo Fisher Scientific) with a Gatan K3 Summit direct electron detector, where each image was composed of 30 individual frames with an exposure time of 3 s and an exposure rate of 16 electrons per second per Å^2^. A total of 4,138 movie stacks were collected. All movie stacks were motion-corrected by MotionCor2 (Zheng *et al.* 2017). Motion-corrected micrographs were imported into cryoSPARC for image processing. The contrast transfer function (CTF) was determined using the “Patch CTF Estimation” option, and the micrographs with “CTF fit < 5 Å” were selected using the “Manually Curate Exposures”. Then 157 particles were manually picked and subjected to 2D classification into 10 classes, 3 of which were selected as the template for the auto-particle picking, yielding 944,822 particles. Several rounds of 2D classification were then performed to remove the poor 2D class averages, and 300,236 particles were obtained. The initial map was built using ab-initio reconstruction without any symmetry applied. Next, three rounds of heterogeneous refinement were performed to further remove bad particles. The final 3D non-uniform refinement was performed using the selected 82,030 particles with or without C3 symmetry applied, and a 3.4-Å map and a 3.7-Å map were obtained, respectively. Resolution for the final maps was estimated with the 0.143 criterion of the Fourier shell correlation curve. Resolution map was calculated in cryoSPARC using the “Local Resolution Estimation” option. The figures were prepared using UCSF Chimera (Pettersen *et al.* 2004) or UCSF Chimera X (Goddard *et al.* 2018).

### Model building

One protomer was first computationally extracted (Pintilie *et al.* 2010) from the 3.4-Å cryo-EM map of the spike protein expressed on HCoV-NL63 coronavirus. The PDB coordinates of biochemically purified HCoV-NL63 spike protein (PDB ID: 5SZS) were then docked into the computationally extracted S protomer map. Residues 883-889 and 993-1000 that were previously unresolved were modeled using SWISS-MODEL (Waterhouse *et al.* 2018). The resultant model was refined using phenix.real_space_refine (Adams *et al.* 2012) and manually optimized with Coot (Emsley *et al.* 2010). The atomic model of protomer was then fitted into the cryo-EM density of the other protomers in the HCoV-NL63 spike trimer using Chimera (Pettersen *et al.* 2004), followed by the optimization of the whole model with phenix.real_space_refine. The glycans in the PDB ID: 5SZS model were also used in our model for refinement. The final model was evaluated by MolProbity (Chen *et al.* 2010) and Q-scores (Pintilie *et al.* 2020). Statistics of the map reconstruction and model optimization are shown in Table S1. All figures were prepared using Chimera (Pettersen *et al.* 2004) or Chimera X (Goddard *et al.* 2018).

## Acknowledgement

We thank Drs. Corey Hecksel and Patrick Mitchell for expert maintenance of Stanford-SLAC Cryo-EM Center and the SLAC National Accelerator Laboratory for supporting conduct of these studies during a university-wide shutdown. This work was supported by the National Institutes of Health grants (P41GM103832, R01AI148382, P01AI120943, R01GM079429, U24GM129564 to W.C.); DOE Office of Science through the National Virtual Biotechnology Laboratory, a consortium of DOE national laboratories focused on response to COVID-19, with funding provided by the Coronavirus CARES Act.

## Author contributions

W.C. and G.S. supervised the study. J.J. prepared the sample. D.C. performed cryo-EM sample preparation. K.Z. and D.C. collected cryo-EM data. K.Z. performed cryo-EM image processing and structure determination; S.L. and K.Z. built and refined the model; K.Z., S.L., G.D.P., D.C., M.F.S, J.J., and W.C. analyzed data. K.Z., S.L., and G.D.P prepared the figures. K.Z., S.L., D.C., J.J, and W.C. wrote the manuscript with input from all other authors.

## Declaration of interests

All authors declare no competing interest.

## Data deposition

Cryo-EM maps of the HCoV-NL63 Spike protein with its associated atomic model have been deposited in the wwPDB OneDep System under EMD accession code XXX and PDB ID code XXX.

---

**Figure S1.**
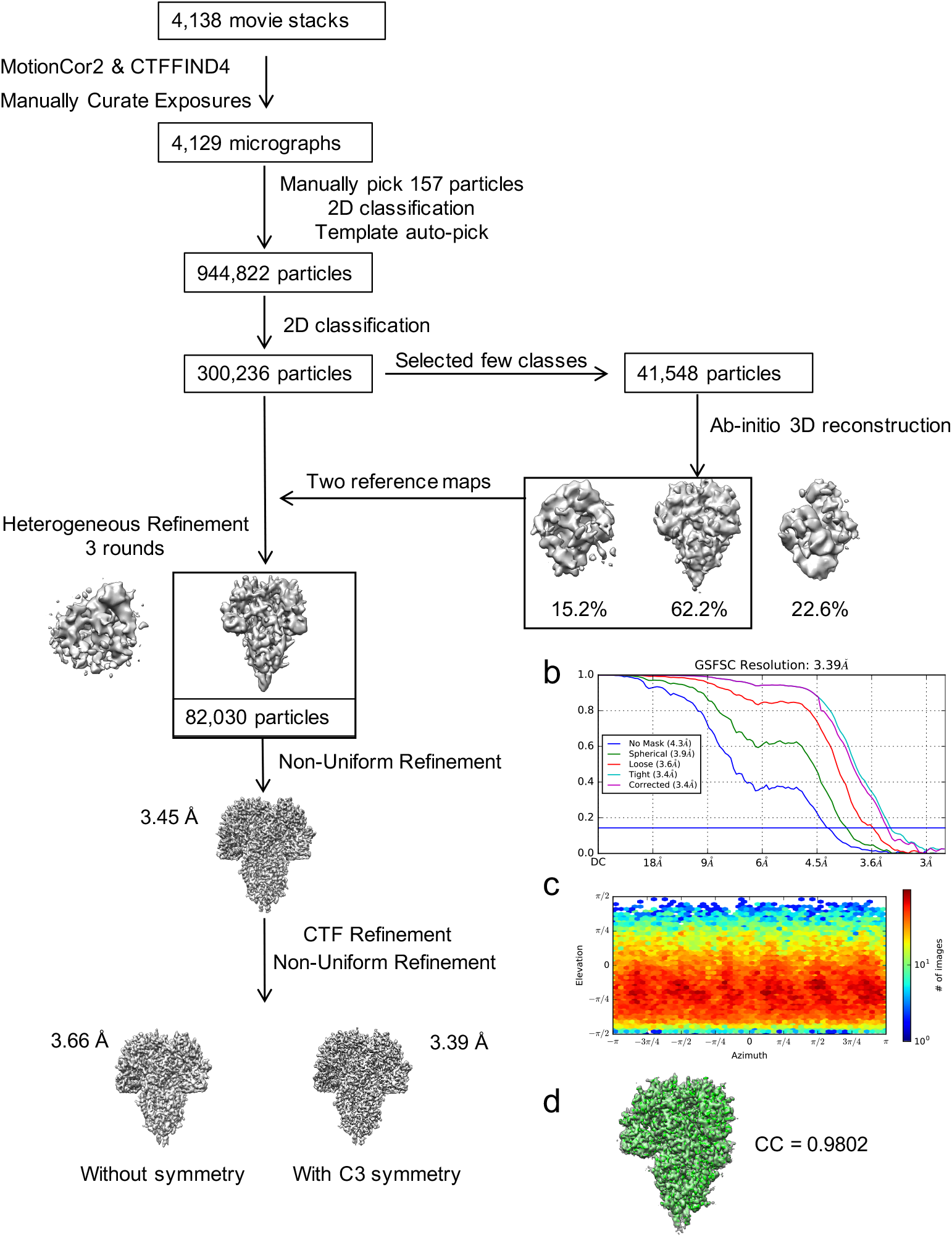
Single-particle cryo-EM data processing of HCoV-NL63 coronavirus spike glycoprotein. (a). Workflow of the data processing. (b) Gold standard FSC plot for the final 3D reconstruction. (c) Euler angle distribution of the particle images of computationally extracted spikes, calculated in cryoSPARC. (d) Cross-correlation coefficient between the 3D reconstructions with (green) and without (grey) C3 symmetry applied.

## Notes

### Competing Interest Statement

The authors have declared no competing interest.

