## Supplementary material for "A 3.4-Å cryo-EM structure of the human coronavirus spike trimer computationally derived from vitrified NL63 virus particles": Table S1

**Cryo-EM data collection, refinement and validation statistics**

|  | HCoV-NL63 S protein  (EMDB-xxxx)  (PDB xxxx) |
| --- | --- |
| **Data collection and processing** |  |
| Magnification | 64k |
| Voltage (kV) | 300 |
| Electron exposure (e–/Å^2^) | 48 |
| Defocus range (μm) | -0.4 - -3.6 |
| Pixel size (Å) | 1.4 |
| Symmetry imposed | C3 |
| Initial particle images (no.) | 944,822 |
| Final particle images (no.) | 82,030 |
| Map resolution (Å)  FSC threshold | 3.39  0.143 |
| Map resolution range (Å) | 2.7-7.0 |
| **Refinement** |  |
| Initial model used (PDB code) | 5SZS |
| Map sharpening *B* factor (Å^2^) | -90 |
| Model composition  Non-hydrogen atoms  Protein residues  Ligands | 30,612  3,576  222 |
| *B* factors (Å^2^)  Protein  Ligand | 89.76  137.32 |
| R.m.s. deviations  Bond lengths (Å)  Bond angles (°) | 0.009  1.214 |
| Validation  MolProbity score  Clashscore  Poor rotamers (%) | 1.44  1.73  0 |
| Ramachandran plot  Favored (%)  Allowed (%)  Disallowed (%) | 91.12  8.80  0.08 |
